# Cell-substrate adhesion drives Scar/WAVE activation and phosphorylation, which controls pseudopod lifetime

**DOI:** 10.1101/732768

**Authors:** Shashi Prakash Singh, Peter A. Thomason, Sergio Lilla, Matthias Schaks, Qing Tang, Bruce L. Goode, Laura M. Machesky, Klemens Rottner, Robert H. Insall

**Author notes:** Correspondence to Robert Insall (. +44/0).

## Abstract

The Scar/WAVE complex is the principal catalyst of pseudopod and lamellipod formation. Here we show that Scar/WAVE’s proline-rich domain is polyphosphorylated after the complex is activated. Treatments that stop activation block phosphorylation in both *Dictyostelium* and mammalian cells. This implies that phosphorylation modulates pseudopods after they have been formed, rather than controlling whether a protrusion is initiated. Unexpectedly, activation-dependent phosphorylation is not promoted by chemotactic signalling, or by signal-dependent kinases such as ERKs, but is greatly stimulated by cell:substrate adhesion. Scar/WAVE that has been mutated to be either unphosphorylatable or phosphomimetic is activated normally, and rescues the phenotype of *scar*^−^ cells, demonstrating that phosphorylation is dispensible for activation and actin regulation. However, pseudopods and patches of Scar/WAVE complex recruitment last substantially longer in unphosphorylatable mutants, altering cell polarisation and the efficiency of migration. We conclude that pseudopod engagement with substratum is more important than extracellular signals at regulating Scar/WAVE’s activity, and that phosphorylation acts as a timer, restricting pseudopod lifetime by promoting Scar/WAVE turnover.

## Introduction

Scar/WAVE is the dominant source of actin protrusions at the edge of migrating cells. In particular, lamellipods (in mammalian cells cultured in 2D) and pseudopods (in cells in 3D environments, or cells such as amoebas) are driven by Scar/WAVE recruiting the Arp2/3 complex, which in turn promotes an increase in the number of polymerizing actin filaments and growth of actin structures (1). It works as part of a large, five-membered complex, whose members have multiple names (2); in this paper they will be referred to as Nap1, PIR121, Scar, Abi and Brk1 in *Dictyostelium*, and Nap1, PIR121, WAVE2, Abi2 and HSPC300 in mammals.

The principal known activator of Scar/WAVE complex activation is the small GTPase Rac. Inactive Rac is GDP-bound, but on stimulation becomes temporarily GTP-bound. The GTP-bound, but not the GDP-bound form binds to the complex (3), in particular through the A-site of PIR121, which includes a Rac-binding DUF1394 domain (4). Interaction with GTP-bound Rac is essential for the complex to be able to function (3, 5). However, though it is clear that Rac is essential, it is not the only regulator (1). Various experiments have found that Rac activation occurs later than the onset of actin-based protrusion (6), and signal-induced actin polymerization can occur earlier than Rac activation (7). In *Dictyostelium*, Scar/WAVE behaviour is much more locally variable than Rac activity (8), so the Rac cannot simply be driving the changes in Scar/WAVE. To understand pseudopod dynamics, it will therefore be vital to enumerate different modes of Scar/WAVE regulation. One potential form of regulation – phosphorylation - has been described in a number of papers, and is also found in untargetted and high-throughput screens. A typical narrative is that Scar/WAVE is phosphorylated in response to external signalling, through kinases such as (particularly) the global signal transducer ERK2. This has been described in cultured fibroblasts (9), MEFs (10), and endothelial cells (11). The phosphorylation is typically found to change the complex from an inactive to an activatable state, so Scar/WAVE phosphorylation directly leads to actin polymerization. Tyrosine kinases, in particular Abl, have been found to be similarly activating (12, 13). These reports are curious, for a number of reasons. First, actin is a strongly acidic protein, so phosphorylation of binding proteins typically weakens their affinity for actin and actin-related proteins. Second, ERK2 has a tightly defined consensus sequence; however, the proposed phosphorylation sites (and confirmed by us below) do not fit this consensus. We have therefore explored the biological functions of Scar/WAVE phosphorylation in detail. A separate set of phosphorylations is present in the C-terminal WCA domain of Scar/WAVE. It is not detectable by, for example, a change in banding pattern on western blots, and difficult to see by mass spectrometry, so is far less widely described. We (14) and others (15) have shown that this is constitutive, and has a role in tuning the sensitivity of the Scar/WAVE complex rather than activating it; both phosphomimetic and unphosphorylatable mutants are active.

One key process in Scar/WAVE biology that is particularly poorly understood is autoactivation. It is clear that pseudopods of migrating cells (which are caused by Scar/WAVE) are controlled through positive feedback – new actin polymerization occurs adjacent to recent pseudopods (8), leading to travelling waves at the edges of cells (16), but the mechanism of this regulation is not well understood (17, 18). It does, however, emphasize the importance of understanding the full dynamics of Scar/WAVE – its recruitment and release from pseudopods, and its synthesis and breakdown – rather than focussing exclusively on its activation.

There is no compelling reason to connect Scar/WAVE phosphorylation to the activation step. Phosphorylation could alter the activity of the complex after it is activated, or alter properties like the rate of autoactivation or the stability of Scar/WAVE once recruited (18). Indeed, in the present work, we find that phosphorylation’s primary role is to control the lifetime of Scar/WAVE after it is recruited to the edge of the cells, meaning that it correlates with pseudopod size rather than the probability of pseudopod generation. We show that phosphorylation occurs after the complex is activated, and at a number of separate, dissimilar sites throughout Scar/WAVE’s polyproline domain. Thus its role appears to be centred around biasing pseudopod behaviour, as required by pseudopod-based models of cell migration, rather than in initiating new pseudopods or actin polymerization.

## Results

### Detailed analysis of Scar/WAVE phosphorylation *in vivo*

A number of authors have observed Scar/WAVE phosphorylation (10, 11, 14, 19, 20). Overall, it is clear that there is constitutive phosphorylation in inactive Scar/WAVE, which we have shown is localised in the extreme C-terminus (14). There is also additional phosphorylation seen in (for example) growth factor stimulated cells, which a number of authors have placed downstream of ERK2 (10, 11, 19), implying it is part of the process by which signalling causes pseudopods to form. To analyse the phosphorylation in detail, we devised an optimised electrophoresis regime. Examined by western blot on normal SDS-PAGE gels, *Dictyostelium* Scar runs as a single band with a diffuse upper edge; quantitative analysis shows this band forms a single, diffuse peak (Figure 1A). To improve separation of different phosphoforms, we optimised PAGE systems. The best separation occurred on 10% acrylamide gels containing low bis-acrylamide concentrations of 0.06% (low-bis gels) compared to the usual 0.3%. In such gels, Scar from migrating *Dictyostelium* resolves into at least six distinct bands (Figure 1B) with clear separation on an intensity plot. Lambda phosphatase resolved the multiple bands to a single one, shifting the multiple intensity peaks (solid line) to single peak (dotted line) (Figure 1C), confirming that the multiple bands are due to phosphorylation, and Scar is typically hyperphosphorylated to varying degrees.

**Figure 1:**
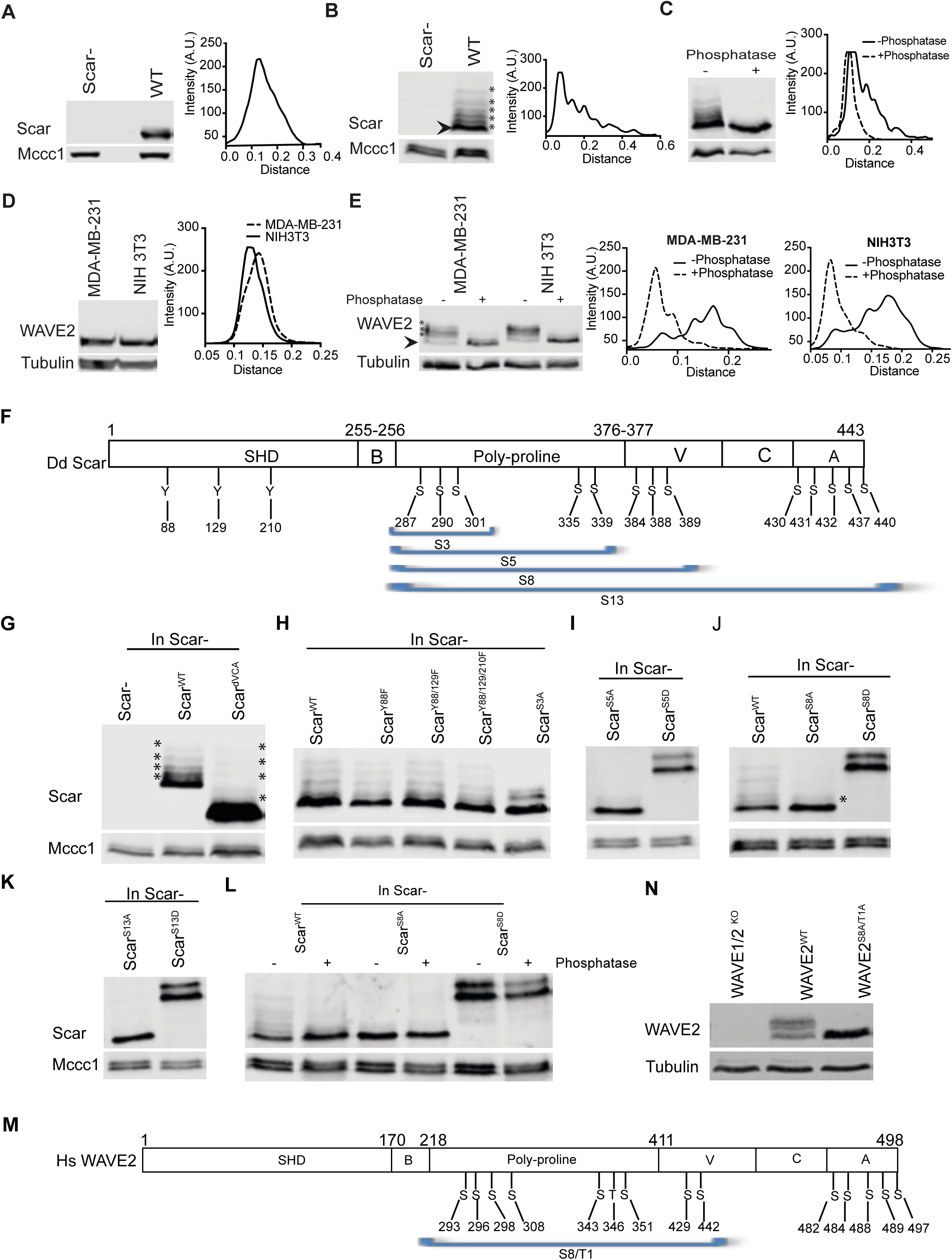
Multiple phosphorylations in Scar/WAVE. **A)** Western blot of *Dictyostelium* Scar using normal (0.3%) bis-acrylamide gels. Whole cells were boiled in sample buffer and immediately separated on 10% bis-tris gels, blotted and probed with an anti-Scar antibody, yielding a single diffuse band. Scan of signal intensity shows a single, slightly diffuse peak. Mccc1 is a loading control (55). **B)** Western blot of *Dictyostelium* Scar using low (0.06%) bis-acrylamide gels. Whole cells were boiled in sample buffer and immediately separated on 10% bis-tris gels, yielding a multiple discrete bands. Scan of signal intensity shows separate peaks. **C)** Phosphatase treatment. Whole *Dictyostelium* cells were lysed in TN/T buffer, then incubated with & without lambda phosphatase, boiled in sample buffer, and analysed using low bis-acrylamide gels. The scan shows multiple peaks resolved into one and shifted downwards after phosphatase treatment. **D)** Western blot of mammalian WAVE2 in mammalian cells using normal (0.3%) bis-acrylamide gels. Whole MDA-MB-231 and NIH 3T3 cells were boiled in sample buffer, separated on 10% bis-tris gels, blotted and probed with an anti-WAVE2 antibody, yielding a single band. Scan of signal intensity shows a single peak. **E)**Western blot of mammalian WAVE2 in mammalian cells using low (0.06%) bis-acrylamide gels. Whole MDA-MB-231 and NIH 3T3 cells were boiled in sample buffer and separated on 10% bis-tris gels, yielding at least 5 bands. Incubation with lambda phosphatase resolves the bands into lower bands. Scans of signal intensity show clear multiple peaks resolving incompletely towards one lower peak. All experiments were repeated thrice with comparable results. **F)** Schematic of *Dictyostelium* Scar architecture and identified phosphorylated residues. **G-K)** Deletions of different amino acids. Lysates from Scar^−^ cells expressing Scar^WT^ and Scar lacking the VCA domain (G), with Tyr 88, 129 & 201 substituted with F (H), Ser 287/290/301 substituted with Ala (H), Ser 287/290/301/335/339 substituted with Ala or Asp (I), Ser 287/290/301/335/339/384/388/389 substituted with Ala or Asp (J), and Ser 287/290/301/335/339/384/388/389/430/431/432/437/440 substituted with Ala or Asp (K), were boiled in sample buffer and analyzed on low (0.06%) bis-acrylamide 10% bis-tris gels. Extra bands in Scar are completely lost after substitution of 13 serines. **L)** Phosphatase treatment of Scar^WT^, Scar^S8A^ and Scar^S8D^. Lambda phosphatase treated cell lysates were analysed using low (0.06%) bis-acrylamide 10% bis-tris gels. Unlike Scar^WT^, Scar^S8D^ shows little removal of additional bands after phosphatase treatment. **M)** Schematic of human WAVE2 architecture and phosphorylated Ser/Thr residues identified in published and high-throughput screens. **N)** Loss of extra WAVE2 bands in B16F1 mouse cells after substitution of polyproline serines. WAVE1/2KO cells were transfected with intact WAVE2 or WAVE2 with Ser 293/296/298/308/343/351/429/442 and Thr 346 substituted with alanine, then lysed, boiled in sample buffer, separated using low (0.06%) bis-acrylamide 10% bis-tris gels, blotted and probed with an anti-WAVE2 antibody.

Similarly, WAVE2 from two mammalian lines (human MDA-MB-231 and mouse NIH3T3) appears on Western blots as a broad band with a single intensity peak (Figure 1D). As with *Dictyostelium* Scar, low-bis gels reveal multiple phosphorylated WAVE2 bands (Figure 1E). Intensity plots show multiple well-resolved peaks that are shifted down after phosphatase treatment confirming the bands represent WAVE2 phosphoforms.

### Identification of phosphorylation sites *in vivo*

Previous reports identified Scar/WAVE phosphosites using protein expressed *in vitro* (11), or overexpressed fusion proteins (9). Because Scar/WAVE’s biological roles occur exclusively as part of the five-membered Scar/WAVE complex, and overexpressed proteins are unevenly and incompletely incorporated into the complex, we analysed the phosphorylation of native proteins. We purified the *Dictyostelium* Scar complex using *nap*1^−^ cells stably transfected with single copies of GFP-Nap1 (14), yielding normal expression levels, and pulled down the complex using GFP-TRAP. Under these conditions Scar is only purified if incorporated in a properly assembled Scar/WAVE complex. Gel-purified Scar bands (Supp. Figure 1A & B) were analysed by LC-MS/MS to identify phosphopeptides. Scar/WAVE is composed of N-terminal SH, central B and polyproline, and C-terminal V, C and A domains (21). We identified three phosphorylated tyrosines (Y88, 129, 210) in the SH domain, three phosphorylated serines (S287, 290, 301) in the polyproline and three (S384, 388, 389) in the V domain (Figure 1F). These are additional to the constitutive phosphorylations found by Ura et al. from 430-440 in the A domain. The number of SCAR forms discernible as bands in western blots from migrating cells is therefore not a surprise – at least 16 sites are phosphorylated *in vivo* in normal cells.

Conspicuously, none of the phosphorylated serines made up MAP kinase consensus sites (Supplementary Table 1). This was surprising, as a number of papers report that Scar/WAVE bands (phosphorylation) are caused by ERK2, which has a strong preference for prolines 2aa N-terminal to and 1aa C-terminal to the target site (22). The sites in human WAVE2 do not fit the MAP kinase consensus (Supplementary Table 1), with the exception of T346, which is relatively sparsely phosphorylated (https://phosphosite.org).

To confirm that we had identified the full range of phosphorylations, we tested which amino acids had to be mutated to block extra bands in westerns. We expressed un-phosphorylatable (Y-F; S-A) and phosphomimetic (S-D) mutants in *scar*^−^ cells. Removing the five phosphorylated serines in the A domain of Scar (14) by deleting the VCA domain (Scar^dVCA^) did not reduce the number of bands (Figure 1G). Mutation of Y88/129/210F in SHD of Scar did not affect the band shifts (Figure 1H). Mutation of the three serines in the polyproline domain (Scar^S3A^, Figure 1H) to alanines reduced the number of bands but did not abolish them. Mutating two more nearby serines (giving Scar^S5A^; S284,290,301,335,339) gave near-complete loss of Scar bands (Figure 1I). Additional mutation of S384/388/389A from the V domain, giving Scar^S8A^ (Figure 1J), yielded Scar that was nearly homogeneous, with a single very faint band above the main Scar population. The residual heterogeneity was lost in cells whose Scar had also lost the sites in the A domain (14), giving Scar^S13A^ (Figure 1I, J & K). Similarly, mutation of all 5 polyproline serines(Scar^S5D^), or all 8 polyproline+V domain serines to aspartate (Scar^S8D^), caused a substantial mobility shift and two distinct bands on low bis gel (Figure 1K). The mobility shift of Scar^S8D^ was also obvious on a normal-bis gel (Supp. Figure 1C), but only a single band of Scar^S8D^ was resolved. These two forms are apparently not caused by phosphorylation; strong phosphatase treatment, which also caused significant proteolysis, caused only partial loss of the upper band (Figure 1L); the difference may be due to a small modifier like ubiquitin, though we were unable to find ubiquitin itself by western blot in any variant (Scar^WT^, Scar^S8A^ or Scar^S8D^; Supp. Figure 2).

The mammalian Scar/WAVE2 has been also shown to be phosphorylated at multiple positions (Figure 1M, Supp. Table 1). Similar to Scar, mutation in WAVE2^S8A/T1A^ (S293/296/298/308/343/351/429/442/T346A) removed all phosphorylated bands (Figure 1N). This confirmed that the multiple bands in WAVE2 are due to S/T phosphorylations.

In conclusion, the multiple bands seen in westerns are mostly caused by differential phosphorylation at sites in the polyproline region, with minimal contributions from the V and A domains.

### Scar/WAVE is phosphorylated after activation

The Scar/WAVE complex must interact with the GTP-bound, active form of Rac1 before it can catalyse Arp2/3 complex activation and pseudopod generation (3, 23–28). We would not expect Rac interactions to be important to the phosphorylation if, as described in several publications (3, 23, 26), the phosphorylation is an upstream process regulating Scar/WAVE activation. We therefore determined the effect of Rac1 binding on Scar/WAVE phosphorylation. First we inhibited the Rac1 activity in both *Dictyostelium* and B16F1 cells by EHT1864, an effective inhibitor of RacGEF activity (29, 30), and assessed the band shifts of Scar/WAVE. EHT1864 treatment resulted in complete loss of Pak-CRIB-mRFPmars2 from cell periphery and depolarized *Dictyostelium* cells (Figure 2A, Suppl. Video 1), confirming that the inhibitor reduced active Rac levels to almost nothing. Similarly, lamellipods of B16F1 cells collapsed after EHT1864 treatment; their F-actin at cell edges also appeared reduced (red, Figure 2B).

**Figure 2:**
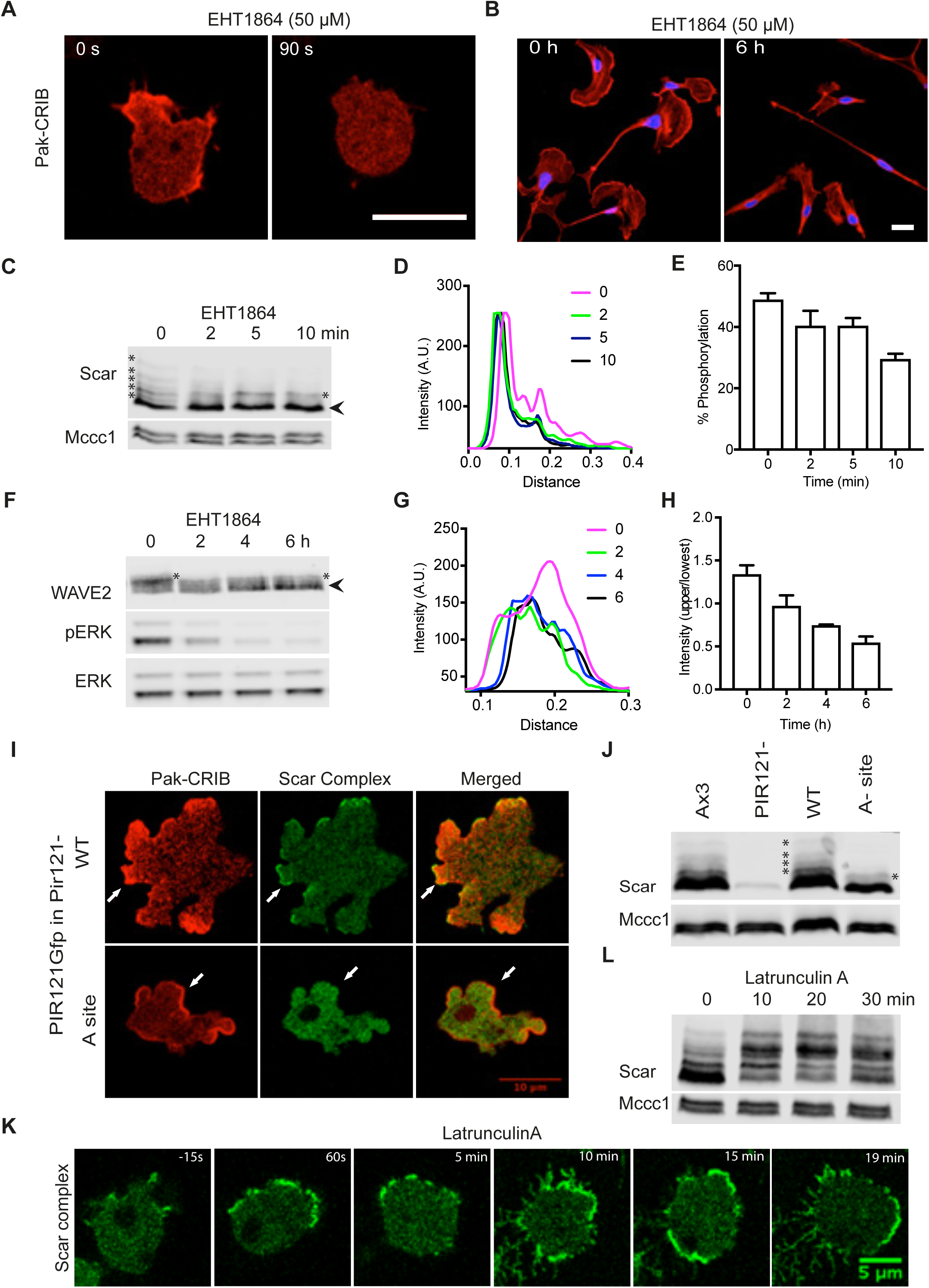
Rac1 binding and Scar/ WAVE phosphorylation. **A)** Effect of Rac1 inhibition by EHT 1864 on Pak-CRIB-mRFPmars2 localization in *Dictyostelium*. *Dictyostelium* cells expressing Rac1 marker, Pak-CRIB-mRFPmars2 were imaged by AiryScan confocal microscopy. Pak-CRIB-mRFPmars2 (Red) rapidly delocalises after addition of EHT1864 (EHT1864, 50 μM) treatment (Scale bar= 10 μm). **B)** Effect of Rac1 inhibition by EHT 1864 on lamellipod formation in B16 melanoma cells. B16F1 cells were seeded on Laminin A coated glass, and treated with EHT1864 (50 μM) for 6h. Cells were fixed and stained with Alexa fluor 568-phalloidin. Untreated cells retained broad lamellipods with intense F-actin at the leading edge, while lamellpods and F-actin are compromised in the EHT1864 treated cells. **C-E)** Effect of Rac1 inhibition by EHT 1864 on *Dictyostelium* Scar phosphorylation. Cells were treated with 50 μM EHT1864 for the indicated times, and lysates were analysed for Scar band shifts by western blotting using low (0.06%) bis-acrylamide gels. Disappearance of extra bands after EHT 1864 treatment indicates loss of phosphorylated Scar. Graph in D shows density quantitation of western blots at different times. E shows ratio of upper (more phosphorylated) band aggregate intensity to total Scar (bars show mean±SEM, n=3). **F-H)** Effect of Rac1 inhibition by EHT 1864 on B16 melanoma WAVE2 phosphorylation. Cells were treated with 50 μM EHT1864 for the indicated times, and lysates were analysed for WAVE2 band shifts by Western blotting using low (0.06%) bis-acrylamide gels. Disappearance of extra bands after EHT 1864 treatment indicates loss of phosphorylated WAVE2. Graph in G shows density quantitation of western blots at different times. H shows ratio of upper (more phosphorylated) band aggregate intensity to total WAVE2 (bars show mean±SEM, n=3). **I)** Requirement for Rac1-binding PIR121 A site for Scar complex localization in *Dictyostelium* cells. Pir121^−^ cells expressing Pir121-eGFP with and without mutated A site (K193D/R194D) were coexpressed with Pak-CRIB-RFPmars2 and imaged by confocal microscopy while migrating under agarose up a folate gradient. Upper panel: unaltered PIR121-eGFP; lower panel: Pir121-eGFP^A site^. Rac1 localizes (Red) to the membrane in both Pir121-EGFP^WT^ and Pir121-EGFP^K193D/R194D^. Scar complex (green) is recruited to sites of Rac1 activation only when Rac can bind. **J)** Requirement for Rac1-binding PIR121 A site for *Dictyostelium* Scar phosphorylation. WT, Pir121^−^ and Pir121^−^ cells expressing Pir121 with and without mutated A site (K193D/R194D) were analysed for WAVE2 band shifts by Western blotting using low (0.06%) bis-acrylamide gels. Phosphorylated Scar bands are absent when PIR121’s A site is mutated. **K)** Localization, distribution and hyperactivation of Scar complex after latrunculin A treatment. *Dictyostelium* cells expressing eGFP-NAP1 were imaged using an AiryScan confocal microscope. 5μM latA was added at t=0. Scale bar = 10 μm. Scar complex activation at the cell membrane increases after addition of Latrunculin. **L)** Increased Scar phosphorylation after latrunculin A treatment. *Dictyostelium* cells were treated with latA for the indicated times, then Scar band shifts were analyzed by Western blotting using low (0.06%) bis-acrylamide gels. Phosphorylated bands become markedly more abundant after latA treatment. All experiments were repeated three times.

To assess the effect of Rac1 inhibition on Scar phosphorylation, *Dictyostelium* cells were incubated with 50µM EHT1864 for 2, 5 and 10 min, lysed, and analysed by low-bis western. Inhibition of Rac1 resulted in a strong reduction in Scar phosphorylation (Figure 2C). The intensity profile shows the disappearance of peaks for polyphosphorylated Scar even more clearly (Figure 2D). Quantification of the fraction of Scar in the upper bands shows approximately 50% reduction (Figure 2E). This figure underestimates the total change in phosphate, because the higher bands that are disproportionately lost contain several phosphates. Similarly, inhibition of Rac1 in B16F1 cells by EHT1864 reduced the intensity of the upper bands of WAVE2 within 2 h; the change became even clearer after 6 h of treatment (Figure 2F). Intensity plots of WAVE2 bands show strong reduction in the highest peak of WAVE2 (0h; pink) after EHT1864 treatment (Figure 2G). Quantification of the ratios of the upper (intense) and lowest bands also suggest a substantial loss of WAVE2 phosphorylation (Figure 2H, again an underestimate of total phosphate loss). This implies that Scar is only phosphorylated after it has bound to Rac1, and thus been activated.

Generalized inhibition of Rac1 activity is strongly likely to cause secondary effects, as many proteins such as kinases (for example PAKs (31)) are important for other aspects of motility and require Rac to be activated. To verify that active Rac1’s effect on Scar phosphorylation is direct, we examined cells in which the Pir121 protein was replaced by Pir121-EGFP, either wild type or the Rac1 non-binding, A-site mutant (3, 5). Pak-CRIB-mRFP-mars2 (8) localizes to the pseudopods and cell periphery in both strains (Figure 2I; Panel I, Suppl. Movie 2), showing that Rac is similarly activated in WT and mutant cells. However, the Scar complex is only activated and localized in cells expressing the functional Pir121EGFP, not the A-site mutant (Figure 2I; Panel II, Suppl. Movie 2). Scar phosphorylation is greatly diminished in the A site mutant (Figure 2J), to a level comparable to that seen in Rac-inhibited cells. This confirms that Scar phosphorylation occurs only after the complex is activated by interaction with Rac1.

To affirm the sequence of activation and phosphorylation, we tested the effect of latrunculin, an inhibitor of actin polymerization (32, 33), that causes exaggerated membrane recruitment of the Scar/WAVE complex in mammalian cells (34) and *Dictyostelium*. In *Dictyostelium*, latrunculin causes concentrated and persistent recruitment of Scar complex to the membrane (Figure 2K, Suppl. Video 3). Upon latrunculin treatment Scar phosphorylation rapidly increases; like the Scar recruitment, the phosphorylation reaches a far higher level than is normally seen (Figure 2L). Thus Scar phosphorylation again correlates with its recruitment and activation, rather than upstream processes.

It is interesting to note that there is currently no biochemical assay for Scar/WAVE activity. The only measure of activation is recruitment to the extreme leading edge, and downstream consequences such as actin polymerization. This phosphorylation assay could therefore be a potentially useful indirect measure of recent Scar/WAVE activation *in vivo*.

### Chemotactic signalling does not control Scar phosphorylation

Earlier reports described the MAP kinase ERK2, induced downstream of chemotactic and growth factor signalling, as the principal driver of Scar/WAVE phosphorylation (9, 11, 19). This is an important issue, because it provides a clear connection between chemoattractant signalling and pseudopod initiation and growth that is otherwise lacking. As described earlier, we questioned this result, because MAP kinase sites are usually restricted to a narrow consensus, whereas the sites we identified were diverse. We therefore examined the effect of extracellular signals and MAP kinases on Scar/WAVE phosphorylation. Growing *Dictyostelium* cells use folate as a chemoattractant. During multicellular development they downregulate folate receptors and express receptors to extracellular cAMP. Treatment of growing cells with folate, or developed cells with cAMP, causes increases in F-actin levels that are thought to be important for chemotaxis (35, 36). To provide a consistent narrative, we examined Scar phosphorylation after both treatments. Despite massive activation of erkB protein, no change in the Scar phosphorylation was seen in either folate- or cAMP-treated cells (Figure 3 A & D), either visually from the gel or when expressed quantitatively as the phosphorylated fraction (Figure 3B, C, E & F).

**Figure 3:**
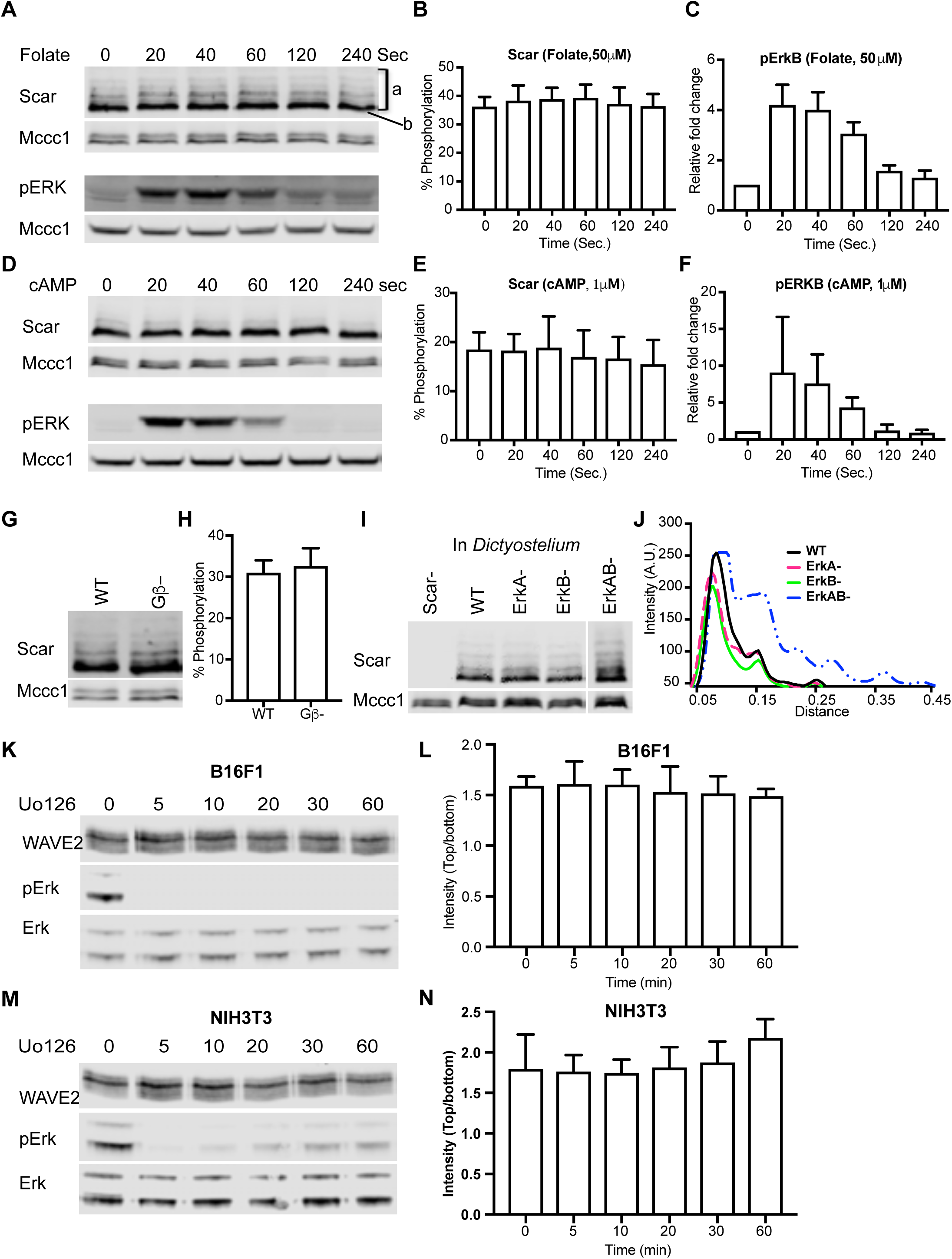
Signalling and Erk independent phosphorylation of Scar/WAVE. **A-C)** Folate stimulation and Scar phosphorylation in *Dictyostelium*. Washed growing cells were treated with 50µM folate for the indicated times, then Scar band shifts were analyzed by Western blotting using low (0.06%) bis-acrylamide gels. Folate does not enhance Scar phosphorylation (B; bands show mean ±SD, n=3) despite substantial signalling response shown by erkB phosphorylation (C). **D-F)** cAMP stimulation and Scar phosphorylation in *Dictyostelium*. Aggregation-stage cells were treated with 50µM folate for the indicated times, then Scar band shifts were analyzed by Western blotting using low (0.06%) bis-acrylamide gels. cAMP does not enhance Scar phosphorylation (E) despite very substantial signalling response shown by erkB phosphorylation (bars show mean±SD; n=3). **G,H)** Scar phosphorylation in WT and Gβ-cells. WT and Gβ-cell band shifts were analyzed by Western blotting using low (0.06%) bis-acrylamide gels. Both show similar band shifts of Scar, as reflected in the quantitation of the upper band (H; bars show mean±SD; n=3) **I)** Scar phosphorylation in MAP kinase (ERK) mutants. Scar^−^, WT, *erk*A^−^, *erk*B^−^ and *erk*AB^−^ cells were analyzed by Western blotting using low (0.06%) bis-acrylamide gels. Single mutants show similar phosphorylation; double mutant shows more, as shown by densitometry of the lanes (J). **K-N)** WAVE2 phosphorylation in B16 (K,L) and NIH3T3 (M,N) cells after inhibition of Erk activation. Cells were treated with U0126 for the indicated times, then analyzed by Western blotting using low (0.06%) bis-acrylamide gels. Erk and phospho-Erk levels were analyzed using normal gels and antibodies against pan-Erk and phospho-Erk. Quantitation of the phosphorylated bands shows no change despite essentially complete inhibition of Erk (bars show mean±SD, n=3).

Knockouts of the *Dictyostelium* Gβ protein show a complete loss of G-protein signalling (37) and reduced ErkB activity (38); since there are also no tyrosine kinase receptors in *Dictyostelium* these cells are complete chemotactic nulls. However, Gβ^−^ mutant cells have normal levels of Scar phosphorylation (Figure 3G & H), as well as apparently normal pseudopods (39). This confirms that chemotactic signalling is not required for Scar phosphorylation. We determined the importance of MAP kinase signalling for Scar/WAVE phosphorylation in both *Dictyostelium* and mammalian cells. First we examined the band shifts for Scar in western blots from *erk*A^−^, *erk*B- and *erk*AB^−^ nulls of *Dictyostelium* (40). The band shifts on the western blot and quantitated peaks were not discernibly different from the WT parent (Figure 3I&J). Since ErkA and ErkB are the only MAP kinases in *Dictyostelium*, MAP kinase signalling is not important for Scar/WAVE phosphorylation. To confirm this result is generally true, we examined the WAVE2 phosphorylation band shifts in mammalian NIH3T3 and B16F1 cells after inhibition of Erk2 activity. Treatment with U0126, an efficient inhibitor of MEK (41) immediately abolished Erk2 phosphorylation (and thus activity) in both NIH3T3 and B16F1 cells, but did not change the phosphorylation state of WAVE2 even after prolonged incubation (Figure 3 K, L, M&N). These results confirm that the Scar/WAVE phosphorylations we observe are neither executed by Erk2 nor do they require Erk2 function.

### Physical adhesion promotes Scar/WAVE phosphorylation

More generally, our data suggest Scar/WAVE phosphorylation is not primarily regulated by extracellular signalling. Another strong candidate regulator of pseudopod and lamellipod is cell:substratum adhesion. In the absence of adhesion, cell protrusions collapse, and the Scar/WAVE complex is strongly localised to the adherent edges of protrusions. We exploited the ability of *Dictyostelium* to grow either in suspension or adhesion. First, we compared the differences in Scar phosphorylation in cells grown in suspension and adhesion in both vegetative and starved cells. The more phosphorylated bands (* in Fig. 4A) are more intense in cells grown adherent to Petri plates, when compared with suspension-grown cells. Similarly, intensity plots of Scar from cells adherent to a surface clearly show more peaks and greater intensities of more-phosphorylated peaks compared with cells grown in suspension (Figure 4B). This observation suggested that adhesion is a strong stimulant of Scar phosphorylation. To investigate this in more detail, we allowed suspensions of cells to adhere to a plastic surface, and observed a clear and highly significant (p= 0.0089; Kruskal Wallis test) upregulation of Scar phosphorylation (Figure 4 C&D). The proportion of intensity in phosphorylated bands increases from 25±3.98% to 42.5±1.99% (Mean±SEM; Figure 4D). Conversely, when we examined adherent cells that had been de-adhered by a stream of buffer from a pipette and kept in suspension, we saw an obvious reduction in phosphorylated bands (Figure 4E). The proportion of intensity in phosphorylated bands decreased from 40±3.59 to 20±3.08 (mean, SEM; Figure 4F). Thus cell:substrate adhesion causes activation of Scar phosphorylation.

**Figure 4:**
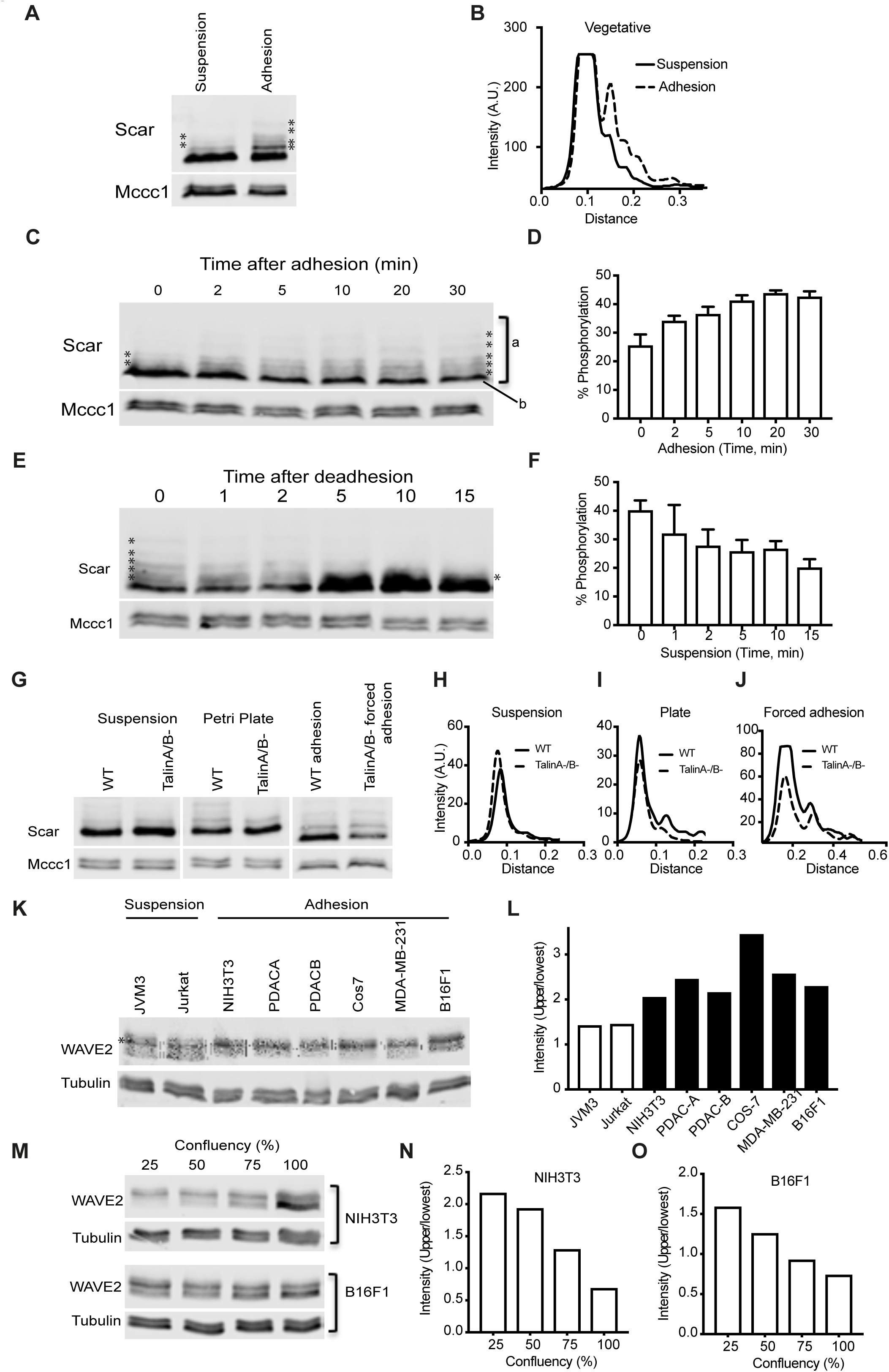
Physical adhesion enhances Scar/ WAVE phosphorylation. **A,B)** Wild type *Dictyostelium* cells were grown in shaken suspension or adhering to Petri dishes, then analyzed by Western blotting using low (0.06%) bis-acrylamide gels. Adherent cells have more and intense bands of Scar/WAVE. Densitometry of lane intensities (mean±SEM, n=3) shows a large difference in the intensities of phosphorylated Scar bands. **C,D)** Acute effect of adhesion on Scar/WAVE phosphorylation. Cells growing in suspension were allowed to adhere in Petri dishes and lysed at indicated time points. Cell lysates were analyzed by Western blotting using low (0.06%) bis-acrylamide gels. The number and intensity of bands (*) increases with time of adhesion. Quantitation of lane intensities (mean±SEM, n=3) confirms a progressive increase in the intensities of phosphorylated Scar bands. **E,F)** Acute effect of de-adhesion on Scar/WAVE phosphorylation. Cells grown in Petri dishes were dislodged by a jet of medium from a P1000 pipette and maintained in suspension (2×10^6^ cells/ml) by shaking at 120 rpm. Cell lysates were analyzed by Western blotting using low (0.06%) bis-acrylamide gels. Quantitation of lane intensities (mean±SEM, n=3) confirms a progressive loss in intensity of phosphorylated Scar bands. **G-J)** Relative requirement of Talin and adhesion. WT and *tal*A^−^/ *tal*B^−^ cells were allowed to make physical contact with plastic Petri dishes, and where appropriate forced to adhere by withdrawing the liquid with 3MM paper. Cell lysates were analyzed by Western blotting using low (0.06%) bis-acrylamide gels. Densitometry of lane intensities shows a large difference between WT and mutant cells on Petri dishes (I), but not when adhesion was forced (J). **K,L)** WAVE2 phosphorylation in adherent and nonadherent mammalian cells. Lysates from cells that grow in suspension (Jurkat and JVM3) and adhesion (NIH3T3, PDAC-A, PDAC-B, Cos7, MDA-MB231 and B16F1) lysates were analysed for WAVE2 band shifts by Western blotting using low (0.06%) bis-acrylamide gels. Suspension-growing cells have additional lower bands (arrow) and less intense phosphorylated (*) band compared to adherent cells. Graph (L) shows the ratio of upper intense band (*) and lowest band of various mammalian cells. **M-O)** Effect of culture density on WAVE2 phosphorylation. NIH3T3 and B16F1 cells were grown at indicated cell densities and lysates were analysed for WAVE2 band shifts by Western blotting using low (0.06%) bis-acrylamide gels. WAVE2 phosphorylation decreases as the cell density increases. * indicates phospho-WAVE2 bands. Graphs show ratio of upper intense band (*) and lowest band. All graphs are representative of three independent experiments.

We examined whether cell:substrate adhesion coupled to Scar activation via adhesion-linked signalling, or through a physical process such as deformation. This was enabled by talin A/B double knockout *Dictyostelium* cells. Talin is a key couple between adhesion molecules like integrins and the actin cytoskeleton (42, 43) and *tal*A/B^−^ mutant cells completely fail to adhere when allowed to fall onto a substratum (44). Consequently they remain extremely spherical and pseudopod-free when cultured, even in dishes where normal cells adhere. However, we were able to force *tal*A/B^−^ cells into adhesion to plastic, glass or 1 % agar surfaces by carefully aspirating the liquid from the dishes so cells were compressed by capillary forces. Scar phosphorylation increases sharply in cells that are making talin-independent contacts with the substratum and the comparison of Scar phosphorylation between suspension and attached *tal*A/B^−^ cells are identical to WT cells (Figure 4G). Intensity plots show no difference in phosphorylation between adherent WT cells and *tal*A/B^−^ cells that have been forced to adhere (Figure 4 H, I & J). This result implies that Scar/WAVE phosphorylation is mediated by the physical process of attachment – through mechanochemical processes, for example - rather than through canonical integrin adhesion signalling. To determine if Scar/WAVE2 phosphorylation in mammalian cells is also adhesion dependent, we compared the band shifts in cells that grow in suspension (Jurkat and JVM3) and adhered to a surface (NIH3T3, the pancreatic cancer lines PDACA & B, Cos7, MDA-MB-231 and B16F1). The Jurkat and JVM3 cells have less intense upper bands, meaning highly phosphorylated Scar/WAVE2 is less abundant. Both also have an additional lower band (representing less-phosphorylated Scar/WAVE2) not seen in any of the cells that grow in adhesion (Figure 4K). The intensity ratios of the upper intense and lowest of the bands present in all cells showed a marked increase in adherent cells (Black bars in Figure 4L) compared with suspension cells (White bars in Figure 4L). We also modulated adhesion by varying cell density – cells grown at high confluency have a lower adhesion area due to space constraints imposed by neighbouring cells. The amount of Scar/WAVE2 phosphorylation decreases with confluency in both NIH3T3 and B16F1 cells (Figure 4M, N & O).

### Unphosphorylated and Phosphomimetic Scar Mutants are Fully Functional

In essentially every lamellipod or pseudopod of every normal cell, actin and Arp2/3 complex recruitment are mediated by the activation of the Scar/WAVE complex (2). Phosphorylation of the polyproline domain has been described as important for Scar/WAVE complex activation and formation of such protrusions (9–11, 14, 19, 20). To determine the importance of polyproline domain phosphorylation to cell migration *in vivo*, we constructed unphosphorylatable (Scar^S8A^) and phospho-mimetic (Scar^S8D^) mutants, with serines mutated to alanines and aspartates, respectively (Figure 1F & J) and expressed them in *scar*^−^ cells. Both mutants were expressed at wild type levels (Figure 1J).

Cells lacking Scar are inefficient in migration, as they have small pseudopods and migrate mainly by blebbing, and are less directional ((8)Figure 5A; Panel I, Suppl. movie 4). In contrast, *scar*^−^ cells rescued with Scar^WT^, Scar^S8A^ and Scar^S8D^ formed longer pseudopods that moved faster and split frequently (Figure 5A; Suppl. Video 4). Pseudopod formation was similar in cells rescued with WT and either Scar mutant. The migration speed of knockout cells was completely rescued by expression of wild type or either phosphomutant Scar (Figure 5B), with phosphorylation changes causing a slight increase in cell speed. The tortuosity of the cell perimeters, which describes the amount of recent protrusion and shape change, was low in *scar*^−^ cells but rescued by Scar^S8A^ and Scar^S8D^ just as well as by Scar^WT^ (Figure 5C). Thus, phosphorylation is clearly neither a precondition for activity, nor a direct cause of SCAR inactivation.

**Figure 5:**
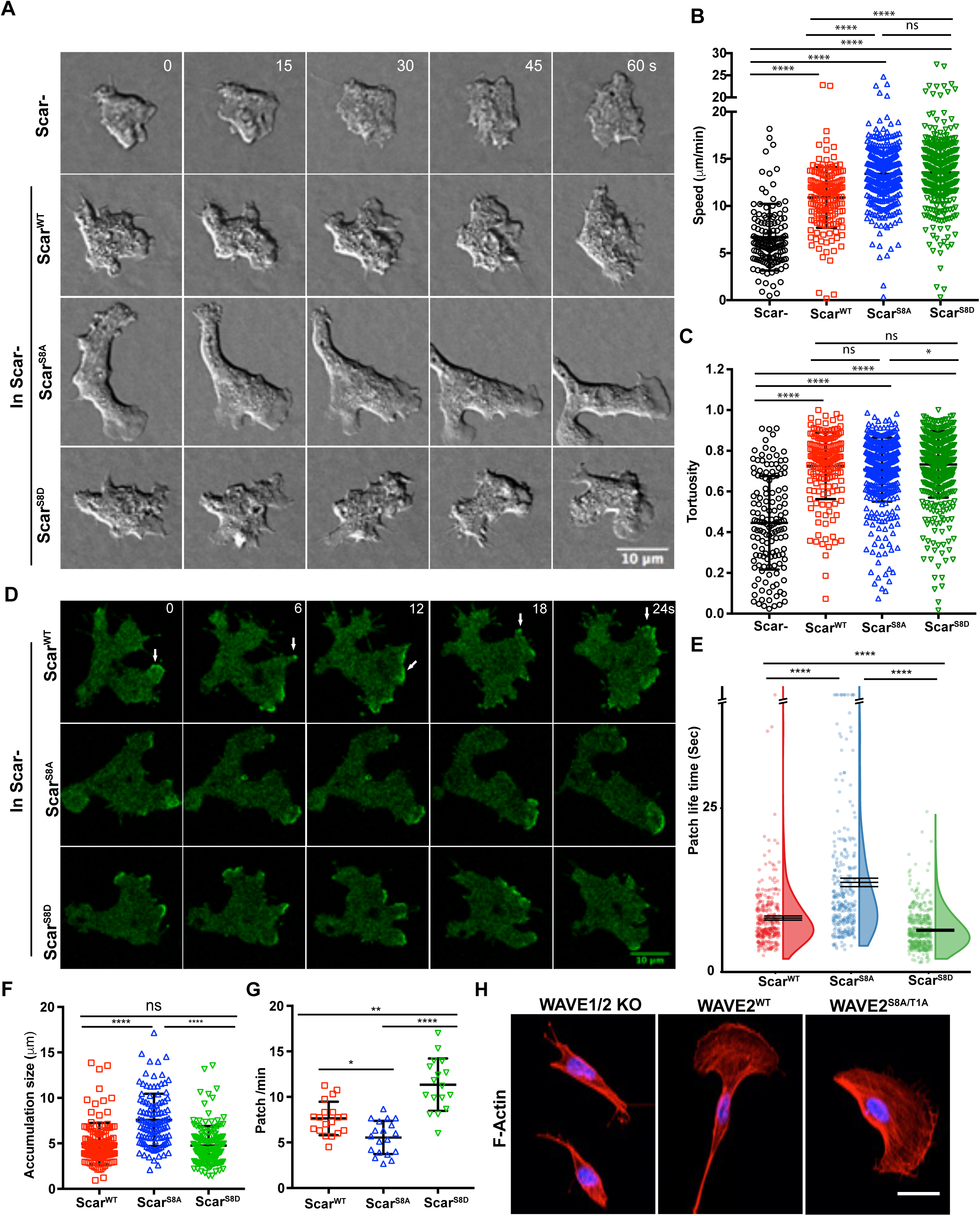
Scar/WAVE phosphorylation mutants are functional. **A-C)** Rescue of pseudopod formation by mutated Scar. Scar-cells were transfected with Scar^WT^, Scar^S8A^ and Scar^S8D^, and allowed to migrate under agarose up a folate gradient while being observed by DIC microscopy. All rescued cells formed actin pseudopods; those formed by Scar^S8A^ expressing cells were broad and elongated. Panel B shows quantitation of speeds and C the tortuosity (total distance migrated/net distance migrated) of paths in B (mean ± SD; n= 140^Scar−^ 170 ^WT^, 413^S8A^, 437^S8D^ over 3 independent experiments; ****p≤0.0001, * p≤0.05, one-way ANOVA, Dunn’s multiple comparison test) **D)** Subcellular localization of Scar complex. Scar^WT^, Scar^S8A^ and Scar^S8D^ were expressed in Scar^−^/Nap1^−^/eGFP-NAP1 cells, allowed to migrate under agarose up a folate gradient, and examined by AiryScan confocal microscopy. All show efficient Scar/WAVE complex (green) localization to the pseudopods. **E,F,G)** Lifetimes, size and generation rates of mutant Scar patches. Scar^−^/Nap1^−^/eGFP-NAP1 cells expressing Scar^WT^, Scar^S8A^ and Scar^S8D^ were allowed to migrate up folate gradients under agarose, EGFP localization was observed by AiryScan confocal microscopy (1f/3s). Patch lifetime was measured by counting the number of frames a patch showed continuous presence of labelled Scar complex, size was measured by hand from single frames, and generation rate calculated from the number of patches lasting at least 2 frames. Scar^S8A^ shows increased and Scar^S8D^ diminished lifetime compared with Scar^WT^. Scar^S8A^ patches are longer-lived than Scar^S8D^ and Scar^WT^, and Scar^S8A^ generates more and Scar^S8D^ less frequently than Scar^WT^ (mean ± SD; n>25 cells; * p≤0.05, ** p≤0.01, **** p≤0.0001, one-way ANOVA, Dunn’s multiple comparison test). **H)** Rescue of lamellipodia formation by WAVE2 phosphomutants. B16F1-WAVE1^−^/2^−^ cells expressing WAVE2^WT^ and WAVE2^S8A/T1A^ were allowed to adhere on laminin A coated coverslips, fixed and stained with Alexa 568-phalloidin. WAVE2^WT^ and WAVE2^S8A/T1A^ both rescue lamellipodia formation.

### Unphosphorylatable Scar is Recruited in Larger and Longer-lived Patches

Activated Scar complex localizes to the fronts of cells and causes actin protrusions there (8). We have found that GFP-tagging Scar itself greatly alters both Scar and pseudopod dynamics. Therefore to examine the effects of Scar phosphorylation on the dynamics of Scar recruitment, we expressed Scar mutants in Scar-cells in which Nap1 has been replaced by a single copy of GFP-Nap1 (i.e. *scar*^−^/Nap1^GFP^ cells; Ura et al., 2012). Scar^WT^, Scar^S8A^ and Scar^S8D^ all localized efficiently at the pseudopod periphery, but there were marked differences in the dynamics of the association (Figure 5D; Suppl. Video 5). As usual (14) Scar^WT^ was recruited in bursts of fairly short duration (8.2 s ± 5.9s, mean±SD). The unphosphorylated Scar (Scar^S8A^) remained recruited for a substantially longer duration (13.7s ± 11.63s, mean±SD). As well as being longer-lived (Figure 5E), Scar^S8A^ patches were also larger (Figure 5F). This led to changes in patch frequency – the larger, longer-lived Scar^S8A^ patches were made at a lower frequency than Scar^WT^ (Figure 5G). Conversely, localization of Scar^S8D^ was even briefer (6.37 ±1.94s, mean±SD) and patches were made at an even higher frequency than Scar^WT^, though they were about the same size. Based on these observations, we hypothesized that Scar/WAVE phosphorylation is not required for its activation, but rather modulates cell migration by controlling the dynamics of Scar/WAVE patches. To confirm this, we constructed nearly completely un-phosphorylated and phosphomimetic Scar by additionally mutating the 5 serine residues of the acidic domain (Ura et al., 2012). Scar complex and F-actin were both beautifully accumulated in the pseudopods of cells expressing Scar^S13D^ and Scar^S13D^ (Suppl. Video 6).

To further confirm this observation in B16 F1 mouse melanoma cells, we employed WAVE1/2 knock out cells and examined the effect of unphospho-mutations in the proline-rich domain (WAVE2S^8A/T1A^; Figure 1N). WAVE1/2 KO cells transfected with empty vector did not form lamellipodia (Figure 5H). However, expressing both WAVE2^WT^ and WAVE2^S8A/T1A^ in knockout cells rescued lamellipodia formation. These results firmly indicate that Scar phosphorylation is neither required for its activation, nor does it cause direct inactivation.

### Scar Phosphomutants Rescue *scar*^−^/*wasp*^−^ Cells

In *Dictyostelium* cells lacking Scar, WASP can be repurposed to drive pseudopod formation and Arp2/3 complex activation (8, 45), which complicates the phenotypes of *scar* mutants. *Scar*^−^/*wasp*^−^ cells cannot move and cannot grow; we have therefore developed a tetracycline-inducible double knockout (*scar*^tet^/*wasp*^−^) cell line to test Scar function without WASP complementation (45). We exploited this line to test unphosphorylatable Scar mutants’ ability to support growth and pseudopod formation. *Scar*^tet^/*wasp*^−^ cells were transfected with Scar^S8A^ and Scar^S8D^ then deprived of doxycycline 48 h prior to experiments to remove native Scar. As usual (46) both mutants express at wild type levels (Figure 6A). Importantly, expression of both Scar^S8A^ and Scar^S8D^ is dominant; expression of the native Scar is suppressed even in the presence of doxycycline (Figure 6A).

**Figure 6:**
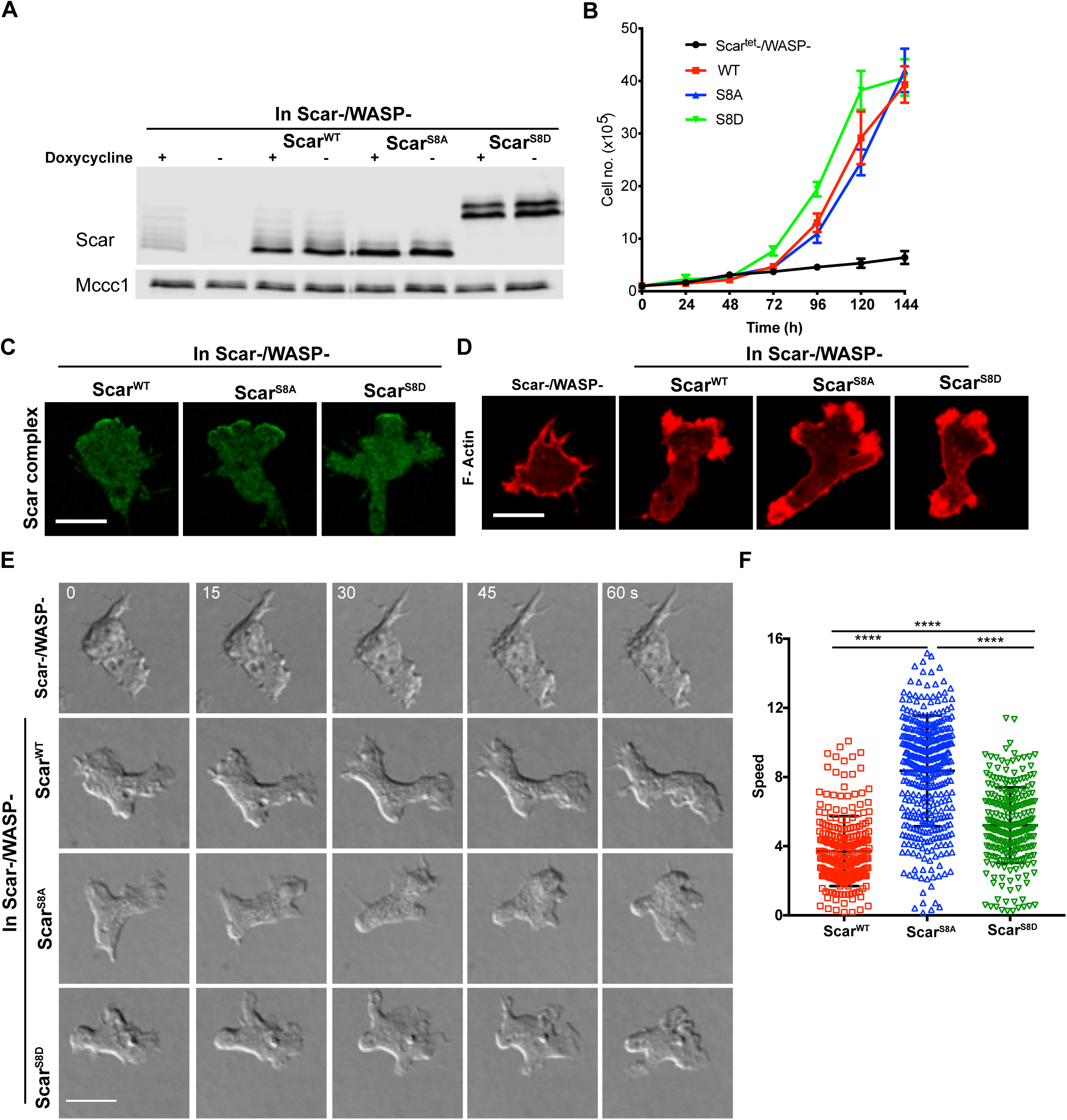
Scar phosphomutants rescue growth and pseudopods in scar^−^ /wasp^−^ cells. **A)** Expression of Scar^WT^, Scar^S8A^ and Scar^S8D^ in cells lacking endogenous Scar and WASP. scar^tet^/wasp^−^ cells expressing Scar^WT^, Scar^S8A^ and Scar^S8D^ were grown with or without doxycycline and analysed by Western blotting using low (0.06%) bis-acrylamide gels. MCCC1 was used as a loading control. **B)** Rescue of Scar/WASP mutant cell growth. scar^tet^/wasp^−^ cells were transfected with Scar^WT^, Scar^S8A^ and Scar^S8D^, expression of endogenous Scar was blocked by removing doxycycline at t=0, and cell growth in suspension was measured using a Casy cell counter (Innovatis). Points show mean±SEM, n=3. **C)** Localisation of Scar complex in scar^tet^/wasp^−^ cells expressing Scar phosphomutants. Scar^WT^, Scar^S8A^ and Scar^S8D^ were co-expressed with HSPC300-EGFP in Scar^tet^/WASP^−^ cells, and cells migrating under agarose up a folate gradient were imaged by AiryScan confocal microsopy. **D)** Localisation of F-actin in scar^tet^/wasp^−^ cells expressing Scar phosphomutants. Scar^WT^, Scar^S8A^ and Scar^S8D^ were co-expressed with LifeAct-mRFPmars2 in scar^tet^/wasp^−^, grown with doxycycline to maintain normal Scar, then kept without doxycyline for 48hrs so only the mutant was expressed. In cells without Scar or WASP, LifeAct was observed in thin spiky filament, but the pseudopods of Scar^WT^, Scar^S8A^ and Scar^S8D^ expressing cells contained normal F-actin levels. **E)** Rescue of pseudopod formation by mutated Scar. scar^tet^/wasp^−^ cells were transfected with Scar^WT^, Scar^S8A^ and Scar^S8D^, and allowed to migrate under agarose up a folate gradient while being observed by DIC microscopy. All rescued cells formed actin pseudopods; those formed by Scar^S8A^ expressing cells were broad and elongated. Scar^WT^, Scar^S8A^ and Scar^S8D^ expressing cells formed pseudopods. Panel F shows quantitation of speeds observed by 10 X phase contrast microscopy). Almost no unrescued scar^tet^/wasp^−^ cells migrated under the agar so no speed could be measured. (mean ± SD; n>278 cells from 3 independent experiments; **** = p≤0.0001, one-way ANOVA, Dunn’s multiple comparison test).

Similar to Scar^WT^, both Scar^S8A^ and Scar^S8D^ rescued the growth of *Scar*^tet^/*wasp*^−^ cells (Figure 6B). To study the migratory phenotype, cells were examined migrating under agarose up a folate gradient. As previously observed (45), repressed *Scar*^tet^/*wasp*^−^ cells made long, stable filopods, but could not form pseudopods or migrate. However, cells expressing either Scar^S8A^ or Scar^S8D^ showed clear localisation of Scar complex to protrusions and F-actin polymerisation (Figure 6 C&D; Suppl. Video 7), and robust pseudopod formation (Figure 6E; Suppl. Video 8). Both mutants rescued cell migration effectively (*scar*^−^/*wasp*^−^ cells do not migrate under the agar, so their speed cannot be measured in this assay, but it is essentially zero). Scar^S8A^ supported a substantially higher speed (8.35 ± 3.2μm/min, mean±SD) than either WT (3.71 ± 2μm/min, mean±SD) or Scar^S8D^ (5.2 ± 2.1 μm/min, mean±SD; Figure 6F). These results firmly indicate that unphosphorylatable and phosphomimetic mutants are fully functional for growth and cell migration in cells without requiring cooperation from WASP.

### Phosphorylation as a Degradation Signal

Scar/WAVE complex dynamics at the pseudopod periphery are poorly understood. In particular, the regulation of Scar/WAVE removal – which must be rapid, to allow the dynamic behaviour observed in moving cells - is mysterious (24). Our results suggest that phosphorylation is among the mechanisms that control the rapid changes in Scar complex localization (Figs 2K, 5 and 6). One way this could be achieved is by promoting breakdown of the complex. This seems particularly appropriate as we previously found that mutations in the acidic domain rendered Scar unstable (14) as well as excessively active. Furthermore, re-examining our earlier data (Figures 4D and 4F, for example) we noticed a surprising change – under conditions where Scar phosphorylation decreases, such as deadhesion, there is an acute increase in the total amount of protein (Figure 7A). Similarly a drop in Scar phosphorylation causes protein levels to increase acutely (Figure 7B). We therefore compared the stability of Scar^WT^, Scar^S8A^ and Scar^S8D^ during starvation, with and without cycloheximide to inhibit protein synthesis.

**Figure 7:**
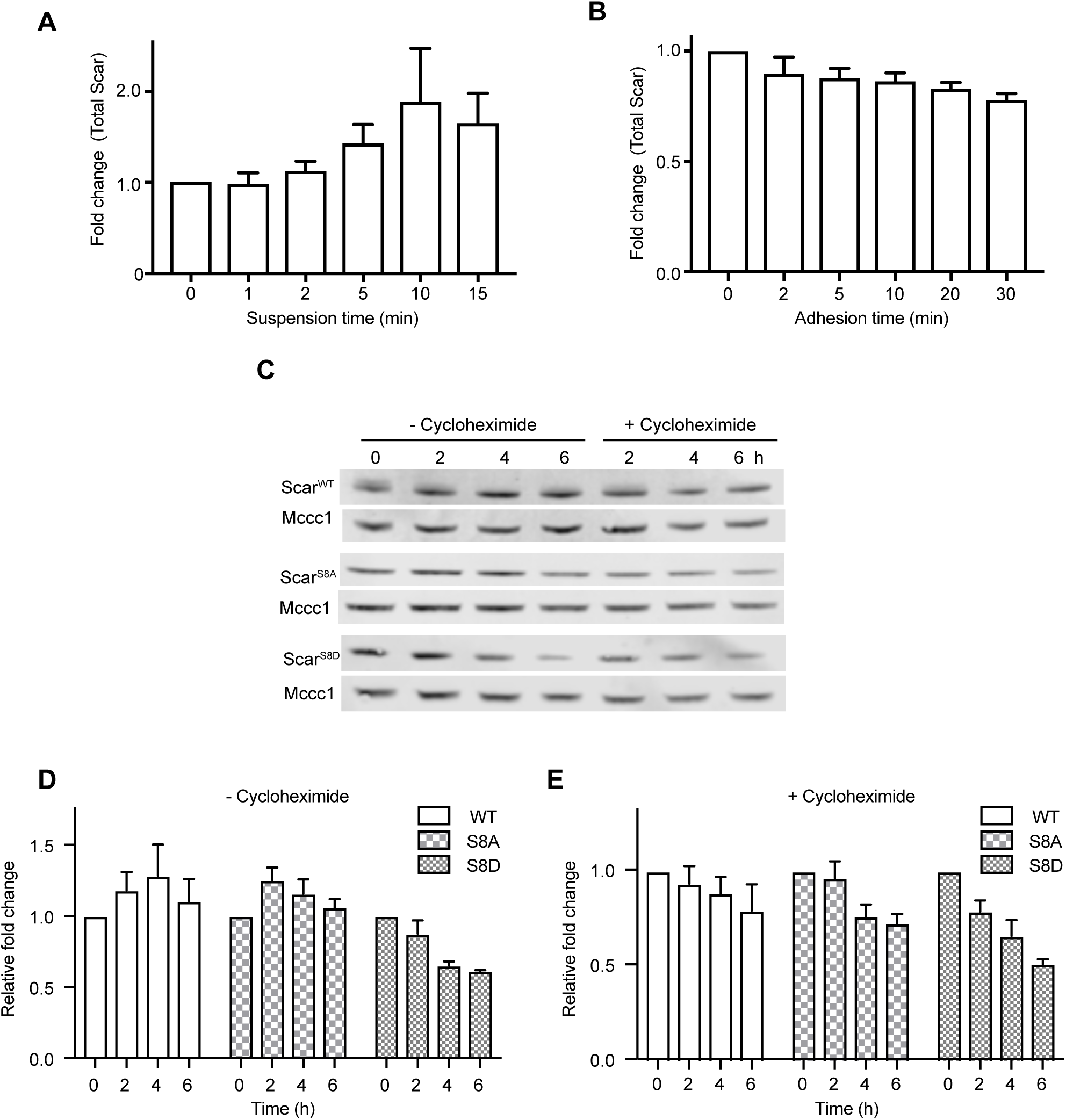
Phosphorylation destabilizes Scar/WAVE. **A)** Total Scar levels from adherent cells dislodged from the surface and maintained in suspension. Data measured by densitometry from cells in Fig. 4F. Deadhesion, which diminishes phosphorylation, acutely increases SCAR levels. **B)** Total Scar levels from nonadherent cells acutely allowed to adhere to a surface. Data measured by densitometry from cells in Fig. 4D. Adhesion, which causes a strong increase in phosphorylation, acutely decreases SCAR levels. **C-E)** Reduced stability of phosphomimetic Scar. Scar^−^ cells expressing Scar^WT^ Scar^S8A^ and Scar^S8D^ were starved with and without cycloheximide. Protein samples were analysed for Scar expression at indicated time points using normal 10% bis-tris gels. The graphs show densitometric quantitation of the Scar bands from panel C, with and without cycloheximide (mean±SEM, n=3).

Starvation makes cells move more rapidly (47) and increases Scar levels (Figure 7 C&D). In the absence of cycloheximide, Scar^WT^ & Scar^S8A^ levels increased as usual during starvation (Figure 7 C&D). Scar^S8D^ levels, however, decreased (Figure 7C&D). When cells were starved with cycloheximide, amounts of Scar^WT^, Scar^S8A^ and Scar^S8D^ all decreased, but the loss was substantially stronger in Scar^S8D^ (Figure 7C & E). This result shows that phosphomimetic Scar is less stable, implying that the reason Scar^S8D^ makes smaller and shorter-lived pseudopods is that it is degraded more rapidly.

## Discussion

We have used an improved assay for phosphorylation of Scar/WAVE’s polyproline domain to open a window on pseudopod dynamics. This has revealed a number of surprises. Phosphorylation occurs on multiple sites after Scar/WAVE is activated by proteins such as activated Rac, and reveals an adhesion-dependent, signalling-independent pathway. It is clear that these phosphorylations are not important for Scar/WAVE to be activated, and it seems likely that they do not inhibit its activation, either. Rather, phosphorylation appears to be a relatively subtle tool by which cells can manipulate the dynamics (both recruitment and loss) of Scar/WAVE patches and thus pseudopods. Pseudopod lifetime intersects with a wide range of cell biology – it underlies the mechanism of chemotactic steering (48, 49), and regulates cell polarity, persistence (50) and shape change (51) – and mutants with uncontrolled pseudopod lifetime have serious motility defects (4). Thus although the function of Scar/WAVE phosphorylation is different from what has been believed, it is fundamentally physiologically important.

Our finding that signalling does not greatly alter the rate of Scar/WAVE phosphorylation, and thus activation, does not imply that signalling is not important to pseudopod formation and evolution. Chemotaxis – cells migrating up gradients of soluble signalling molecules – clearly happens, and is clearly mediated by pseudopods. However, we (48) and others (49) have found that normal chemotaxis rarely involves initiation of new pseudopods – rather, pseudopod generation is mostly random, but localised receptor activation modulate the positions where pseudopods evolve, their lifetime and stability after they are formed, and sometimes the pattern in which new split pseudopods are directed. None of these measured processes require extracellular signals causing direct activation of pseudopod catalysts; rather, more subtle changes like the precise subcellular localisation and timing of Scar/WAVE activation could be biased by chemoattractant signalling. Equally, chemoattractants could work through pathways other than Scar/WAVE activation, biasing the growth and lifetimes of pseudopods rather than their initiation. This is in full agreement with the pseudopod-centred view of chemotaxis (52), which emphasizes that pseudopods are self-organized, rather than the outcome of external signals.

The kinase sites around the polyproline domain have little in common with one another, and do not match the consensus sequences for most kinases that are active in the actin domain. We hypothesize that the phosphorylation sites are sterically inaccessible until the Scar/WAVE complex is activated. Once the complex is active it adopts an open conformation, whereupon any and all of a large range of constitutive kinases such as CK2 and GSK3 could act on it. In this case the phosphorylation would be a passive reflection of the fact that the complex had been activated, rather than a driver or even modulator of the initial activation process. This explanation is comfortably consistent with the wide range of data we have obtained, though it paints a different picture of Scar/WAVE activation from what we had expected.

This work showcases our ability to use *scar*^−^/*wasp*^−^ cells to test the functioning of mutated Scar proteins, and the phenotypic changes caused by alterations in the sequence. Previously, results have been complicated because WASP changes its organization and cell biological role when the Scar complex is not present (8). We have therefore been working against an ill-defined background of partial rescue. In this work we show that either phosphomimetic or unphosphorylatable Scar is fully able to rescue the mutant phenotypes, because *scar*^−^/*wasp*^−^ *Dictyostelium* cells cannot grow, do not make pseudopods, and barely move. Similarly, the use of WAVE1/WAVE2 knockout B16-F1 cells allows detailed study of physiologically normal levels of WAVEs without partial complementation. We therefore look forward to a new focus on physiological function and a more direct connection between the dynamics of the Scar/WAVE complex, the behaviour of pseudopods and lamellipods, and cell migration.

Our results imply that cell:substrate adhesion is a crucial driver of Scar/WAVE activation. Adhesion has been an undeservedly neglected area of Scar/WAVE biology. It seems intuitively obvious that pseudopod formation should be regulated by adhesion – if pseudopods or lamellipods are generated in a location where adhesion is difficult or impossible, they will be inefficient or ineffective at producing cell migration. Cells should in retrospect therefore be expected to monitor adhesion, and to favour pseudopods that can attach. The results in Figure 4 showing normal Scar/WAVE activation in talin knockouts offer one reason why connections have so far been missed – activation does not require the integrin-linked signalling pathways that dominate adhesion of (for example) mammalian fibroblasts (53). Instead, we hypothesize that the connection is physical, by direct mechanical coupling. This is a well-documented and common mechanism for coupling proteins and cell behaviour (54). Overall, this work exposes a new and conserved mechanism for regulating migration, which will influence cells moving in many physiological contexts. We look forward to seeing new biological examples emerge.

## Acknowledgements

We thank Prof. Robert R. Kay for erkA and erkB knockout cells, Dr. Rachana Patel for providing NIH3T3 cells, Drs Margaret O’Prey, Heather Spence and Jamie Whitelaw for help in microscopy and mammalian cell culture, and Dr. Simona Buracco for discussions about protein stability.

## Authors’ contribution

RI and SPS conceived and designed experiments. SPS performed experiments. PT assisted SPS during in experiments and provided essential reagents. SL performed MS. SPS wrote the manuscript. RI and LM corrected and improved the manuscript. MS, QT, BG and KR generated B16-F1 mutant cell lines and plasmids

## Supplementary Figure Legends

**Supplementary Figure 1:**
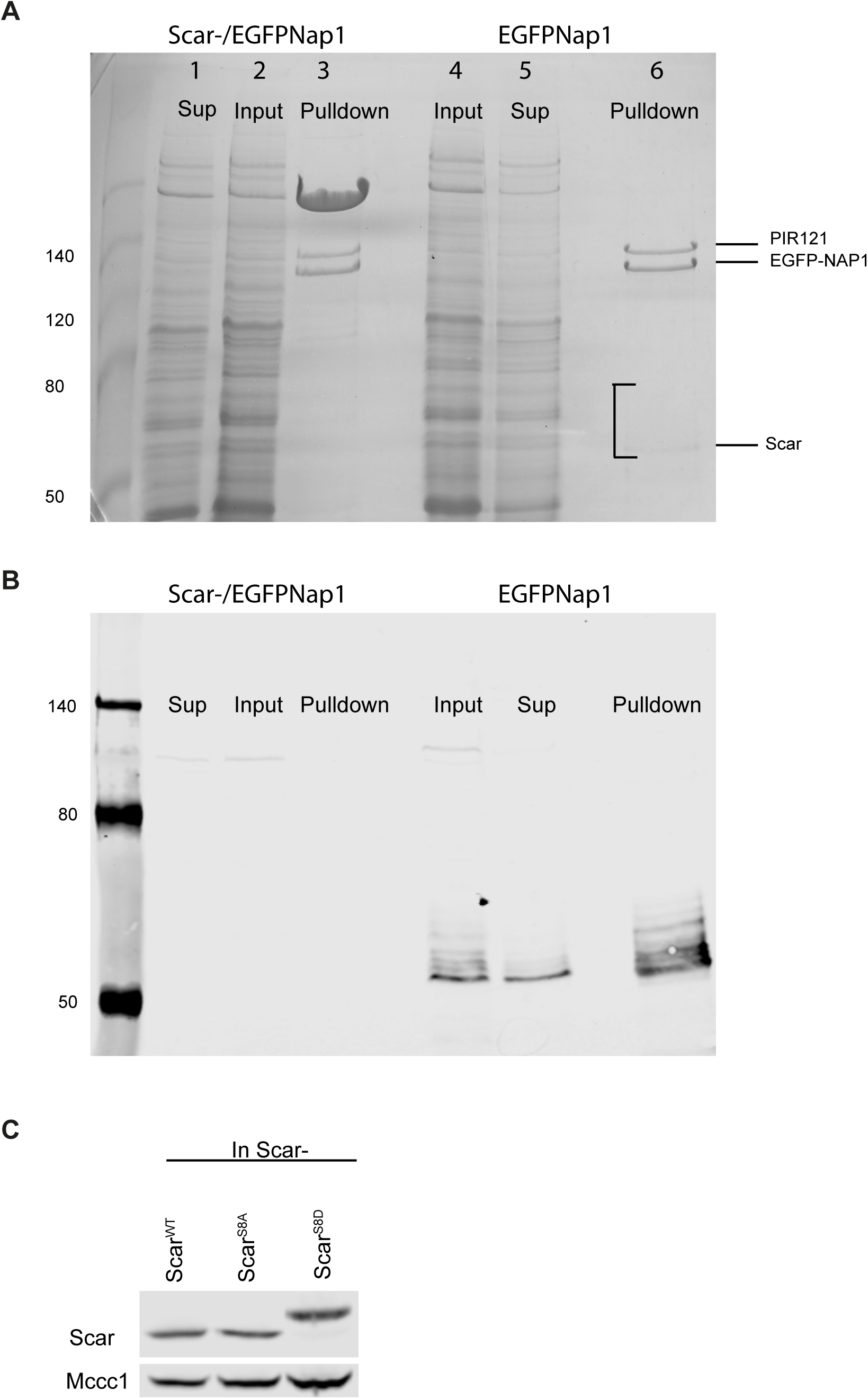
GFP-TRAP pull down of EGFP-NAP1 cells. A) Coomassie brilliant blue stained low-bis acrylamide PAGE gel of GFP-TRAP pull down samples. Lane 1,2,3 indicates samples from Scar-/EGFPNAP1 and lane 4, 5, 6 indicates Samples from EGFPNap1 cells. The Scar band indicated on the gel was exiced for LC-MS/MS. B) Representative western blot of above gel indicating phosphorylated Scar bands.

**Supplementary Figure 2:**
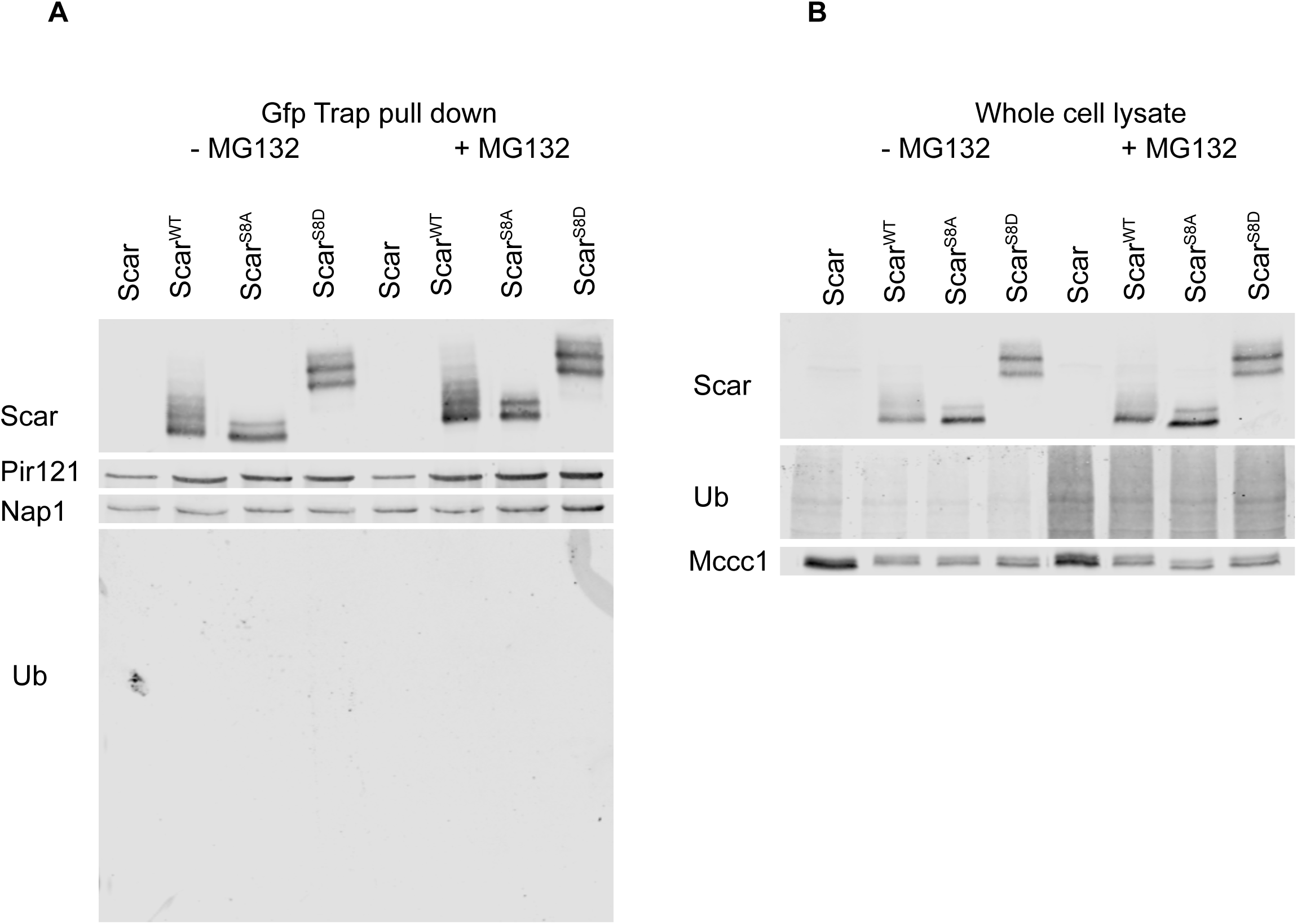
Incorporation of Scar mutants in the complex. A) GFP-TRAP pull down was performed using MG132 treated and untreated *Dictyostelium* Scar-/EGFP-NAP1 cells expressing Scar^WT^, Scar^S8A^ and Scar^S8D^. The eluates were analysed on low-bis PAGE western blotting for Pir121/Nap1/Scar and ubiquitin. B) Scar expression in the cell lysates was analysed by low-bis PAGE and western blotting.

**Video 1. Effect of EHT1864 on Rac1 and Scar complex localization.** *Dictyostelium* cells expressing PakCRIB-mRFPmars2 were allowed to migrate under agarose up folate gradient and observed by AiryScan confocal microscopy. Filmed at 1 frame/2s, Movie shows 10 frames/sec. EHT1864 was added at frame 7 (after14 secs) in the video.

**Video 2: Scar/WAVE and Rac1 activation in cells with mutant PIR121 A site.** Pir121 knock out cells expressing WT Pir121-EGFP and A site Pir121-EGFP were further expressed with PakCRIB-mRFPmars2. Scar complex (green) and PakCRIB-mRFPmars2 (Red) localization was visualized in migrating cells under agarose up folate gradient. Filmed at 1 frame/2s, Movie shows 10 frames/sec.

**Video 3: Effect of Latrunculin treatment on Scar complex localization.**

eGFP-NAP1 cells were seeded on LabTekII rcover glass chambers and imaged by AiryScan imaging. 5µm LatrunculinA was added to the cells undergoing imaging. Filmed at 1 frame/15s, Movie shows 10 frames/sec. Latrunculin was added at the frame 2 (after 15 sec).

**Video 4: Pseudopod formation in Scar phosphomutants.**

Scar^−^ cells expressing Scar^WT^, Scar^S8A^ and Scar^S8D^ were allowed to migrate under agarose up a folate gradient and observed by DIC. Filmed at 1 frame/2s, Movie shows 10 frames/sec.

**Video 5: Scar complex localization in Scar phosphomutants.**

Scar-/EGFP-Nap1 cells expressing Scar^WT^, Scar^S8A^ and Scar^S8D^ were allowed to migrate under agarose up folate gradient and Scar complex activation in pseudopods were observed by airyScan confocal microscopy. Filmed at 1 frame/2s, Movie shows 10 frames/sec.

**Video 6: Scar complex activation in total Scar phosphomutants.**

Scar-/EGFP-Nap1 cells expressing Scar^S13A^ and Scar^S13D^ were allowed to migrate under agarose up folate gradient and Scar complex activation in pseudopods were observed by airyScan confocal microscopy. Filmed at 1 frame/2s, Movie shows 10 frames/sec.

**Video 7: Scar complex activation in Scar^−^/wasp^−^ cells expressing phosphomutant Scar.**

Scar^tet^/wasp^−^ cells expressing Scar^WT^, Scar^S8A^ and Scar^S8D^ were allowed to migrate under agarose up folate gradient and Scar complex activation in pseudopods were observed by AiryScan confocal microscopy. Filmed at 1 frame/2s, Movie shows 10 frames/sec.

**Video 8: Pseudopod formation in Scar^−^/wasp^−^ cells expressing phosphomutant Scar.**

Scar^tet^/wasp^−^ cells expressing Scar^WT^, Scar^S8A^ and Scar^S8D^ were allowed to migrate under agarose up folate gradient and were observed by differential interference contrast microscopy. Filmed at 1 frame/2s, Movie shows 10 frames/sec.

---

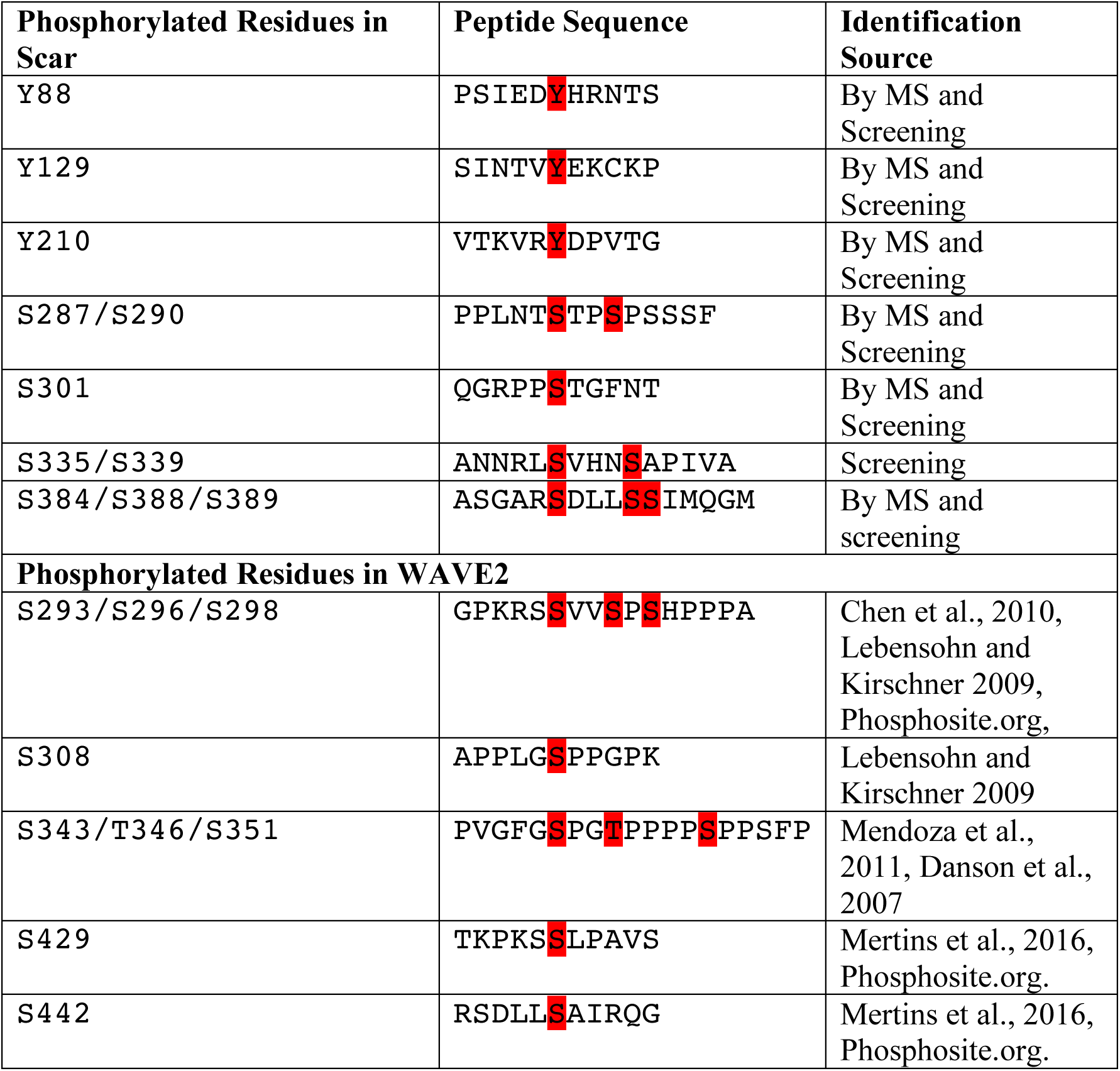

